# TcZC3HTTP, a regulatory element that contributes to *Trypanosoma cruzi* cell proliferation

**DOI:** 10.1101/2023.07.17.549355

**Authors:** Bruno Accioly Alves Romagnoli, Aline Castro Rodrigues Lucena, Eden Ribeiro Freire, Isadora Filipaki Munhoz da Rocha, Lysangela Ronalte. Alves, Samuel Goldenberg

## Abstract

Post-transcriptional regulation of gene expression is a critical process for adapting and surviving *Trypanosoma cruzi*, a parasite with a complex life cycle. RNA binding proteins (RBPs) are key players in this regulation, forming ribonucleoprotein complexes (mRNPs) and RNA granules that control transcript stability, localization, degradation, and translation modulation. Understanding the specific roles of individual RBPs is crucial for unraveling the details of this regulatory network. In this study, we generated null mutants of the TcZC3HTTP gene, a specific RBP in the Trypanosoma family, characterized by a C3H zinc finger and a DNAJ domain associated with RNA and protein binding, respectively. Through cell growth assays, we demonstrated that the absence of TcZC3HTTP or the expression of an additional tagged version significantly impacted epimastigote growth, indicating its contribution to cell proliferation. TcZC3HTTP was found to associate with mRNAs involved in cell cycle and division in epimastigotes, while nutritionally stressed parasites exhibited associations with mRNAs coding for other RBPs and rRNA. Furthermore, our analysis of TcZC3HTTP protein partners revealed the presence of several enzymes during normal growth conditions, whereas starvation conditions enriched ribosomal proteins and other RBPs. This study provides insights into the post-transcriptional regulation of gene expression in *T. cruzi*, highlighting the role of TcZC3HTTP as an RBP involved in cell proliferation and uncovering its versatile functions in different cellular contexts.

**Importance:** Understanding how *Trypanosoma cruzi*, the causative agent of Chagas disease, regulates gene expression is crucial for developing targeted interventions. In this study, we investigated the role of TcZC3HTTP, an RNA binding protein, in post-transcriptional regulation. Our findings demonstrate that TcZC3HTTP is essential for the growth and proliferation of epimastigotes, a stage of the parasite’s life cycle. We identified its associations with specific mRNAs involved in cell cycle and division and its interactions with enzymes and other RBPs under normal and starvation conditions. These insights shed light on the regulatory network underlying gene expression in *T. cruzi* and reveal the multifaceted functions of RBPs in this parasite. Such knowledge enhances our understanding of the parasite’s biology and opens avenues for developing novel therapeutic strategies targeting post-transcriptional gene regulation in *T. cruzi*.

## INTRODUCTION

Over several decades, extensive research has established that *Trypanosoma cruzi* and other trypanosomatids primarily rely on post-transcriptional mechanisms to regulate gene expression (1–4). These mechanisms encompass various events related to RNA metabolism, including RNA maturation, nuclear exportation, subcellular localization, translation, and degradation/storage (5, 6).

Messenger RNAs (mRNAs) undergo individual or coordinated regulation within the cell through dynamic interactions with RNA-Binding Proteins (RBPs). These interactions lead to the formation of ribonucleoprotein complexes (RNPs). Furthermore, RNPs can interact with and exchange components between them, giving rise to larger regulatory complexes known as RNA granules (7). Interactions involving RBPs and their target RNAs are highly dynamic and influenced by environmental changes such as temperature, nutrient availability, and oxidative stress. *T. cruzi* encounters these challenges during its transition within and between mammalian and triatomine hosts (8). The parasite must undergo a coordinated and rapid cellular/molecular response to maintain homeostasis and ensure survival, which includes rearrangements and modulation of RNP complexes. These rearrangements directly impact gene expression. Since RBPs are involved in every step of RNA metabolism, they play a pivotal role as crucial regulators of gene expression (9, 10).

Besides coordinating the cellular RNA levels, many studies in Trypanosomatids that managed to silence or knockout RBPs reported alterations in cellular proliferation, division, and differentiation, indicating that these regulatory proteins also impact major cellular processes (11–17). In *Trypanosoma brucei*, for example, the RBP TbPUF9 silencing decreased bloodstream-form cell growth and increased the number of cells accumulated into the G2/M phase (11). For *T. cruzi*, TcZC3H31 knockout abolished the differentiation into metacyclic trypomastigotes (16), whereas TcZC3H12 disruption presented a higher metacyclogenesis rate and a decrease in cellular proliferation (18).

The bacterial adaptive immune system CRISPR/Cas9 discovery, and its use as a genome engineering tool, emerged as a promising alternative to perform gene disruption in several organisms, including *T. cruzi* (19–21). Recently, our group developed an approach that allowed us to confirm the first knockout achieved with CRISPR/Cas9 of an RBP in *T. cruzi*, the C3H zinc finger protein TcZC3HTTP, thus unveiling a new perspective to advance in RBPs characterization in this organism (22).

Here we report our findings regarding the disruption of TcZC3HTTP in *T. cruzi,* an RBP exclusive to the Trypanosoma family. Modulating TcZC3HTTP expression, we identify that this RBP is related to *T. cruzi* cell proliferation. Next, we evaluated the mRNA targets and protein partners of TcZC3HTTP in normal and nutritional stress conditions. We also performed immunoprecipitation assays followed by RNA-seq and mass spectrometry using a protein-tagged version (TcZC3HTTP-3xFLAG). In addition, we assessed total RNA and protein in these conditions to investigate the effects on the TcZC3HTTP mRNA target and protein partner candidates in the *tczc3*http null mutant population.

## RESULTS

### TcZC3HTTP phylogeny

To understand the TcZC3HTTP role in *T. cruzi* biology, we first investigated when this protein emerged in the Trypanosoma family history. Throughout phylogenetic analysis performed with databases from different trypanosomatids combined with sequence alignments, we were able to find TcZC3HTTP orthologs in most Trypanosoma species, all presenting the zinc finger C3H and DNAJ domains (Figure 1 and Supplementary Figure S1). Interestingly, this RBP is absent in two unrelated groups within this family, the subfamily Strigomonadinae and a branch inside the subfamily Trypanosomatinae that includes the species *T. evansi*, *T. equiperdum*, *T. congolense*, and *T. brucei*. Since all the other species, which consists of the entire subfamilies Leishmaniinae, Blechomonadinae, *Phytomonas sp*, and the remaining Trypanosomatinae representatives along with *T. cruzi*, presented the gene *tczc3http*, it likely appeared in the Trypanosoma common ancestor. It is unclear, however, if this RBP was lost on two different occasions or if its appearance occurred only after the Paratrypanosomatinae group speciation and then lost in the *T. brucei* and related Trypanosomatinae species group. However, it is possible to assume that, besides being old, the relationship of TcZC3HTTP with Trypanosomatids seems unique since this protein was found exclusively in this group.

**Figure 1.**
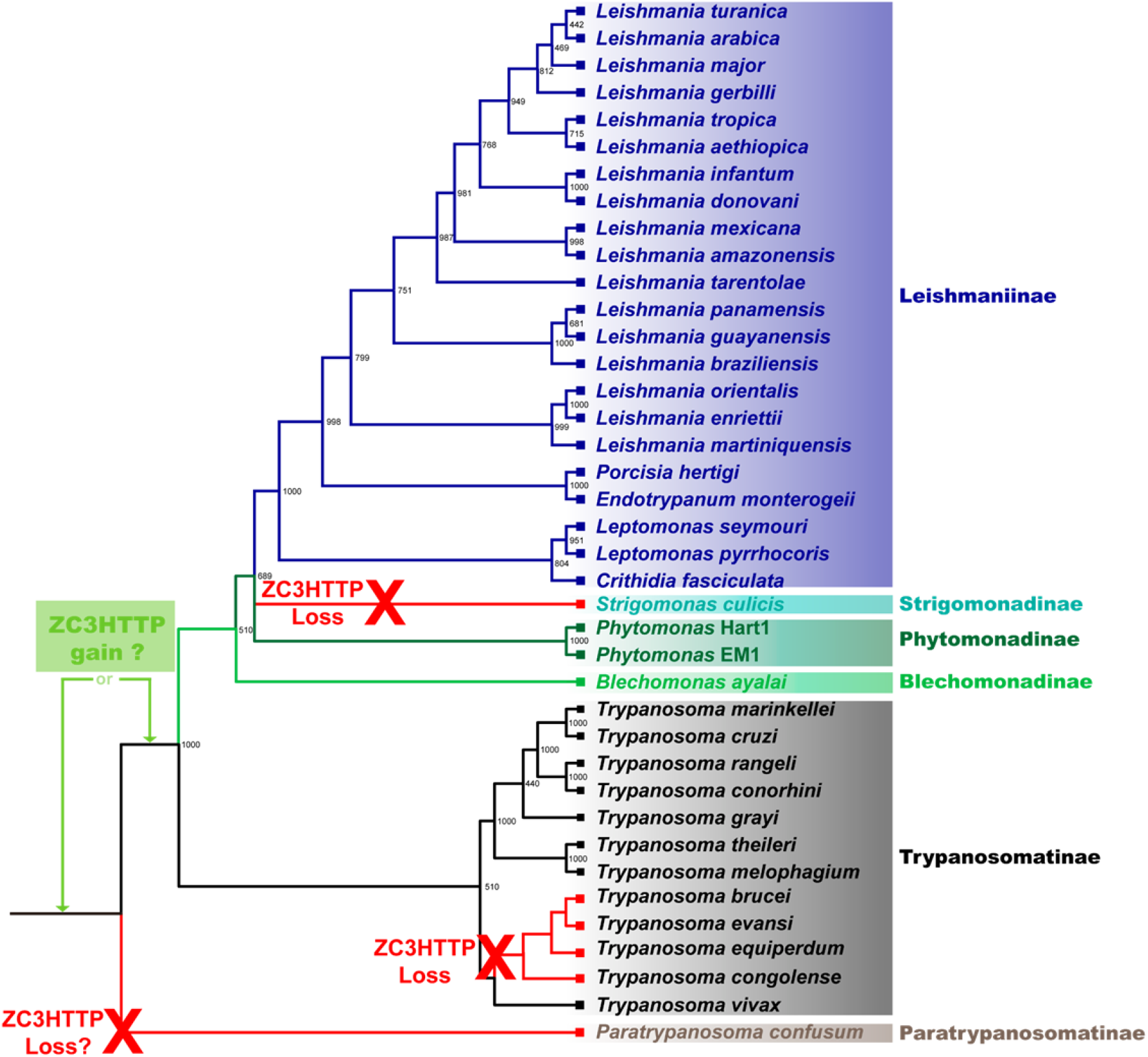
Maximum likelihood tree of Trypanosomatids ZC3HTTP proteins. The tree was inferred using the Maximum Likelihood method based on PhyML analysis for the best evolutionary model. Bootstrap values are shown at the node branches. The tree is drawn to scale, with branch lengths measured in the number of substitutions per site. Colors indicate ZC3HTTP sequences for Trypanosomatids subfamily groups.

### Impact of TcZC3HTTP on gene expression regulation in *T. cruzi*

In our study, we generated two distinct populations of *T. cruzi* to investigate the role of TcZC3HTTP. The first population, designated as ΔTcZC3HTTP, lacked the TcZC3HTTP gene, while the second population expressed a tagged version of TcZC3HTTP known as TcZC3HTTP-3xFLAG [22]. We employed the CRISPR/Cas9 system with two specific guide RNAs (gRNA 99 or gRNA 231) to disrupt the TcZC3HTTP gene in the ΔTcZC3HTTP population, and successful knockout was confirmed by sequencing analysis [22].

Subsequently, we examined the growth characteristics of these populations. The results revealed a significant growth defect in the ΔTcZC3HTTP population compared to the control group, evident from the third day of the growth curve and persisting until the seventh day (Figure 2A). Conversely, the population expressing the tagged version of TcZC3HTTP (TcZC3HTTP-3xFLAG) exhibited a higher growth rate than the control group (Figure 2B). The cellular localization of the protein was also evaluated in epimastigotes under normal growth conditions and epimastigotes subjected to nutritional stress. It is possible to observe a slight granular pattern in stressed parasites compared to unstressed epimastigotes (Figure 2 C).

**Figure 2.**
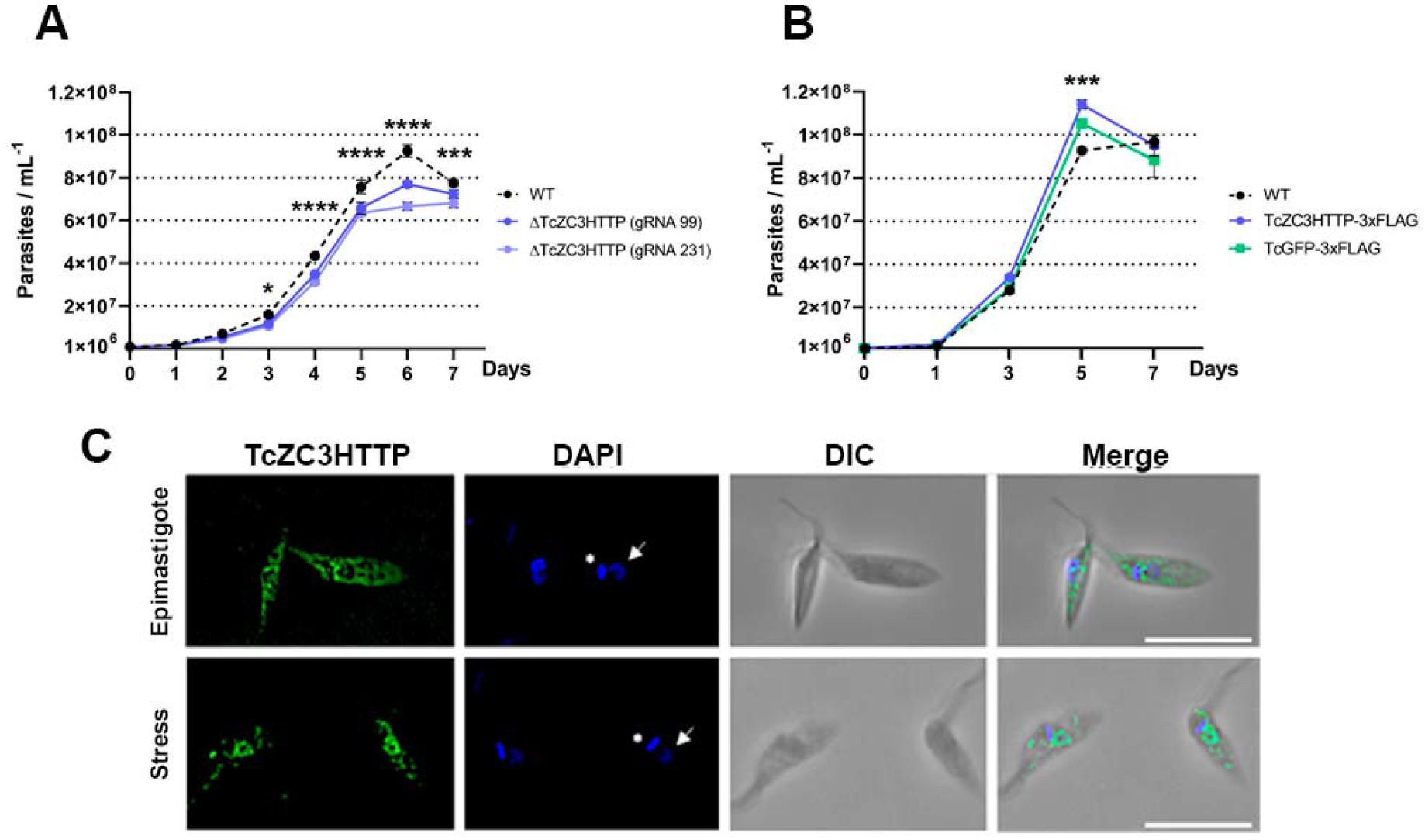
TcZC3HTTP disruption (A) or TcZC3HTTP-3xFLAG expression (B) impact. Growth curve showing cellular density (y-axis) throughout cultivation days (x-axis) for null mutants for TcZC3HTTP or parasites expressing TcZC3HTTP-3xFLAG. Wild-type (WT) and parasites expressing eGFP-3xFLAG (TcGFP-3xFLAG) were used as control (n=3; *p<0,05;***p<0,001;****p<0,0001). All graphs and statistical analyses were made within the GraphPad Prism software. (**C**) Cellular localization of the TcZF3HTTP-flag protein in epimastigotes under normal growth conditions and parasites subjected to nutritional stress. The commercial monoclonal antibody α-FLAG® (1:1000) was used. Nucleus DNA (white arrow) and kinetoplast (asterisk) were stained with DAPI, and the obtained images were overlaid with DIC (merged). The scale bar represents 10 µm.

These results suggest that TcZC3HTTP plays a role in regulating cell proliferation in *T. cruzi*. The absence of TcZC3HTTP led to decreased growth, while the presence of the tagged protein enhanced cellular proliferation. These observations highlight the significance of TcZC3HTTP in controlling *T. cruzi* cell proliferation and provide valuable insights into its potential functions in the parasite’s life cycle and development.

### TcZC3HTTP RNA targets

To elucidate the role of TcZC3HTTP in gene expression regulation in *T. cruzi*, we analyzed the RNAs pulled down with the protein-tagged version of TcZC3HTTP under two conditions: epimastigotes and nutritionally stressed parasites (Figure 3). Our investigation revealed 243 transcripts associated with TcZC3HTTP-3xFLAG in normal conditions and 159 in stress conditions (Supplementary Tables 1 and 2). Among these transcripts, 214 were exclusively found in epimastigotes, 130 in stressed parasites, and 29 in both conditions.

**Figure 3.**
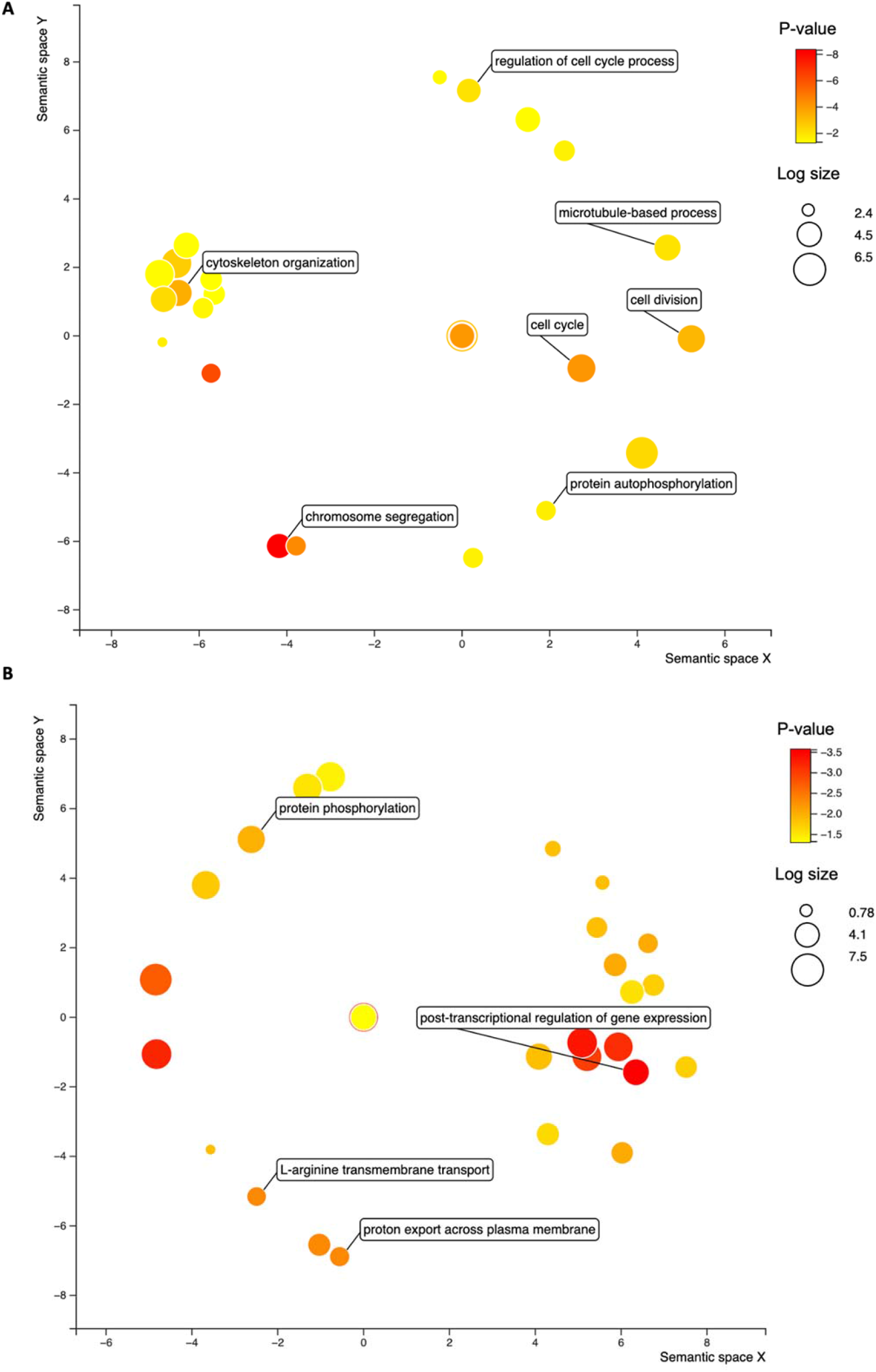
Bubble chart with the gene ontology terms enriched in the RNAs associated with TcZC3HTTP in epimastigotes (**A**) and stressed parasites (**B**). The colors refer to the p-value (log-10), and the bubble size the term counts (in log scale). The distance of the bubbles in the x- and y-axis refer to the proximity of the terms that belong to close-related biological processes.

Upon querying the Tritryp database for information on these transcripts, we discovered that 40% of the enriched transcripts (148 out of 373) encoded hypothetical proteins. We conducted individual searches for domain predictions and functional indications to gain further insights into these transcripts. The resulting information allowed us to summarize the transcripts that encoded known products associated with TcZC3HTTP mRNP, as depicted in Table 1.

**Table 1.** Top transcripts associated with TcZC3HTTP in epimastigotes*.

| TritrypDB ID | Identification | Log<sub>2</sub> FC | FDR p-value |
| --- | --- | --- | --- |
| TcCLB.503453.20 | TcZC3HTTP | 5,82 | 0,0% |
| TcCLB.503869.40 | TcPUF9 | 2,62 | 0,0% |
| TcCLB.507521.110 | TcPUF5 | 1,86 | 0,9% |
| TcCLB.511815.50 | Zinc finger protein family member | 2,09 | 0,2% |
| TcCLB.510295.15 | ATP-dependent DEAD/H RNA helicase | 2,09 | 2,4% |
| TcCLB.506679.140 | 40S ribosomal protein S10 | 2,16 | 0,0% |
| TcCLB.503395.50 | 40S ribosomal protein S12 | 1,84 | 0,5% |
| TcCLB.511527.34 | 60S ribosomal protein L8 | 2,07 | 0,1% |
| TcCLB.511163.40 | Cell cycle sequence binding phosphoprotein (RBP45) | 3,25 | 0,1% |
| TcCLB.506711.30 | Cyclin 3 | 2,61 | 5,1% |
| TcCLB.511025.120 | Cyclin 6 | 2,51 | 0,0% |
| TcCLB.509507.49 | Cyclin 8 | 1,89 | 1,3% |
| TcCLB.506583.40 | Cell division related protein kinase 3 (TcCRK3) | 2,52 | 2,4% |
| TcCLB.505183.140 | Cell division protein kinase | 1,93 | 0,2% |
| TcCLB.508917.10 | Polo-like protein kinase (PLK) | 2,74 | 0,0% |
| TcCLB.507713.20 | WD domain, G-beta repeat | 2,86 | 0,3% |
| TcCLB.506327.50 | Centrosomal protein of 57 kDa | 2,77 | 1,5% |
| TcCLB.508919.30 | Centrosomal protein of 120 kDa | 3,18 | 0,0% |
| TcCLB.508317.20 | Centrosomal protein of 162 kDa | 3,70 | 1,0% |
| TcCLB.506529.590 | Centrosomal protein of 164 kDa | 2,59 | 0,0% |
| TcCLB.509717.40 | Centrosomal protein POC5 | 2,06 | 0,5% |
| TcCLB.510101.480 | Centriole protein POC11 | 2,33 | 0,2% |
| TcCLB.507641.190 | Kinetoplastid kinetochore protein 1 (KKT1) | 2,13 | 0,1% |
| TcCLB.511575.70 | Kinetoplastid kinetochore protein 4 (KKT4) | 1,87 | 0,6% |
| TcCLB.509429.140 | Kinetoplastid kinetochore protein 6 (KKT6) | 2,54 | 0,2% |
| TcCLB.506925.490 | Kinetoplastid kinetochore protein 7 (KKT7) | 2,66 | 0,0% |
| TcCLB.507895.110 | Kinetoplastid kinetochore protein 8 (KKT8) | 3,59 | 0,0% |
| TcCLB.508543.80 | Kinetoplastid kinetochore protein 9 (KKT9) | 4,39 | 0,1% |
| TcCLB.509027.60 | Kinetoplastid kinetochore protein 10 (KKT10) | 1,70 | 0,5% |
| TcCLB.511807.120 | Kinetoplastid kinetochore protein 11 (KKT11) | 3,32 | 0,0% |
| TcCLB.505071.50 | Kinetoplastid kinetochore protein 12 (KKT12) | 2,03 | 2,8% |
| TcCLB.507681.250 | Kinetoplastid kinetochore protein 14 (KKT14) | 2,59 | 0,1% |
| TcCLB.506295.30 | Kinetoplastid kinetochore protein 24 (KKT24) | 1,91 | 1,1% |
| TcCLB.507221.50 | Kinetoplastid kinetochore protein 25 (KKT25) | 3,17 | 0,0% |
| TcCLB.419417.19 | Kinetochore interacting protein 1 (KKIP1) | 4,44 | 1,8% |
| TcCLB.510055.70 | Kinetochore interacting protein 5 (KKIP5) | 2,71 | 1,5% |
| TcCLB.509353.50 | Nucleus and spindle associated protein 2 (NuSAP2) | 2,11 | 0,1% |

\*Transcripts related to RNA metabolism are shadowed in green, associated with the cell cycle in blue, centrosome in yellow and kinetochore in orange

In epimastigotes, TcZC3HTTP is associated with mRNAs mainly related to kinetochore proteins, cyclins, and centrosomal proteins, and this result reinforces the role of this protein in cell proliferation. We also identified the cell cycle sequence binding phosphoprotein RBP45 and regulatory elements like the RBPs TcPUF5 and TcPUF9 (Table 1). Accordingly, gene ontology (GO) analysis showed an enrichment of terms involving cell cycle (cyclins), cytoskeleton (centriole), and transcription (Figure 3A).

Interestingly, we observed that the transcripts with the most significant fold change for stressed parasites were ribosomal RNAs (rRNAs) (Supplementary Table S2). Among the mRNAs, the one coding the enzyme phosphoenolpyruvate carboxykinase (PEPCK) exhibited remarkable enrichment (fold change > 120) in TcZC3HTTP-3xFLAG immunoprecipitation during nutritional stress. Also, alpha and beta tubulin transcripts were highly represented, with fold changes exceeding 35 and 45, respectively. Furthermore, we discovered that mRNAs encoding the RBPs TcZC3H11, TcZC3H35, TcZC3H28, and TcPUF6 were enriched (Table 2). The significant gene ontology (GO) terms associated with this condition were primarily linked to microtubules, ATP binding, zinc finger C3H, and coiled-coil domains (Figure 3B).

**Table 2.** Top transcripts associated with TcZC3HTTP in stressed epimastigotes*.

| TritrypDB ID | Identification | Log<sub>2</sub> FC | FDR p-value |
| --- | --- | --- | --- |
| TcCLB.457295.10 | 5.8S ribosomal RNA (M3) | 8,78 | 0,0% |
| TcCLB.419325.10 | ribosomal RNA small subunit | 9,53 | 0,0% |
| TcCLB.473013.10 | ribosomal RNA small subunit | 9,48 | 0,0% |
| TcCLB.467341.10 | ribosomal RNA small subunit | 8,87 | 0,0% |
| TcCLB.445323.10 | ribosomal RNA small subunit | 8,16 | 0,0% |
| TcCLB.447665.10 | ribosomal RNA large subunit alpha | 9,01 | 0,0% |
| TcCLB.509005.91 | ribosomal RNA large subunit beta | 8,90 | 0,0% |
| TcCLB.411483.20 | ribosomal RNA large subunit gamma (M1) | 10,71 | 0,0% |
| TcCLB.422723.20 | ribosomal RNA large subunit delta (M2) | 8,46 | 0,0% |
| TcCLB.509005.121 | ribosomal RNA large subunit epsilon (M4) | 11,65 | 0,0% |
| TcCLB.508355.260 | 60S acidic ribosomal protein (P0) | 5,29 | 0,6% |
| TcCLB.503395.40 | 60S ribosomal protein L18 | 3,76 | 1,0% |
| TcCLB.508823.120 | ribosomal protein S20 | 5,66 | 0,3% |
| TcCLB.509671.70 | ribosomal protein L15 | 3,96 | 4,9% |
| TcCLB.506925.120 | eukaryotic initiation factor 5a (EIF5A) | 3,72 | 5,4% |
| TcCLB.510963.90 | elongation factor 2 | 4,37 | 0,0% |
| TcCLB.508461.140 | TcPABP2 | 4,05 | 4,6% |
| TcCLB.511285.120 | ATP-dependent RNA helicase (HEL67) | 4,26 | 3,3% |
| TcCLB.503453.20 | TcZC3HTTP | 2,75 | 5,1% |
| TcCLB.510729.220 | TcZC3H28 | 2,61 | 2,7% |
| TcCLB.511267.24 | TcZC3H35 | 4,17 | 3,6% |
| TcCLB.504929.5 | TcZC3H11 | 3,78 | 5,1% |
| TcCLB.506945.70 | MKT1 | 4,29 | 3,4% |
| TcCLB.510125.10 | TcPUF6 | 3,08 | 3,4% |
| TcCLB.511025.120 | Cyclin 6 | 3,24 | 2,9% |
| TcCLB.511907.260 | cell division cycle protein | 4,75 | 0,0% |
| TcCLB.510357.10 | J-binding protein (JBP1) | 2,22 | 3,9% |
| TcCLB.508441.20 | glycosomal phosphoenolpyruvate carboxykinase (PEPCK) | 6,95 | 0,0% |
| TcCLB.411235.9 | alpha tubulin | 5,15 | 0,0% |
| TcCLB.506563.40 | beta tubulin | 5,51 | 0,0% |
| TcCLB.507765.20 | PUF nine target 1 | 4,78 | 1,9% |
| TcCLB.511753.130 | U3 small nuclear ribonucleoprotein (snRNP) | 4,15 | 3,7% |
| TcCLB.507021.20 | ESAG8-associated protein | 3,89 | 5,1% |
| TcCLB.509247.30 | CCR4-NOT transcription complex subunit 1 | 3,05 | 0,2% |

\* In green transcripts related to RNA metabolism, in blue cell cycle, and yellow protein folding.

In both experimental conditions, we successfully detected the presence of TcZC3HTTP’s mRNA. Remarkably, we observed a significant enrichment of its transcript in epimastigotes, with a fold change exceeding 50, making it the fourth most abundant RNA identified. Furthermore, TcZC3HTTP mRNA exhibited significant presence even under stress conditions, with a fold change greater than 6. This finding strongly suggests that TcZC3HTTP is involved in an autoregulatory mechanism.

### TcZC3HTTP Protein partners

Given the potential for RBPs to interact with other proteins and the presence of a protein-protein interaction domain (DNAJ domain) in TcZC3HTTP, we used the tagged version of TcZC3HTTP to investigate its candidate protein partners through immunoprecipitation. Following filtering steps, we successfully identified 181 and 55 proteins as modulated under normal and stress conditions, respectively (Supplementary Tables S3 and S4). Notably, we observed a high enrichment of TcZC3HTTP in our experiment, with fold changes of 133 in epimastigotes and 82.1 in stressed parasites, confirming the efficacy of the immunoprecipitation approach.

Proteomic analysis of the TcZC3HTTP immunoprecipitation revealed the presence of numerous proteins, including hydrogenases, kinases, reductases, peptidases, and tRNA synthetases (Tables 3 and 4). Additionally, we identified chaperones, retrotransposon hot spot proteins (RHS), and proteins associated with the proteasome, translation initiation, and elongation factors in epimastigotes. Notably, we observed the association of TcZC3HTTP with several metabolic enzymes, which have been previously recognized as moonlighting proteins (Table 3). This intriguing finding suggests potential functional diversification and the multifaceted roles of TcZC3HTTP in cellular processes.

**Table 3.** Proteins associated with TcZ3HTTP-3xFLAG in epimastigote by immunoprecipitation*.

| TritrypDB ID | Identification | FC | p-value |
| --- | --- | --- | --- |
| TcCLB.503453.20 | TcZC3HTTP | 132.99 | 3.6E-04 |
| TcCLB.506241.170 | 40S ribosomal protein S6 | 7.91 | 2.2E-02 |
| TcCLB.511163.40 | cell cycle sequence binding phosphoprotein (RBP45) | 11.58 | 7.0E-03 |
| TcCLB.510163.20 | elongation factor 1-gamma (EF-1-gamma) | 5.71 | 3.1E-03 |
| TcCLB.511277.70 | Elongation factor Tu, mitochondrial | 8.35 | 7.3E-03 |
| TcCLB.506925.130 | eukaryotic translation initiation factor 5A | 25.48 | 8.0E-04 |
| TcCLB.511277.290 | aconitase | 9.12 | 2.7E-05 |
| TcCLB.510945.70 | aspartate aminotransferase | 25.54 | 3.0E-05 |
| TcCLB.511277.110 | citrate synthase | 6.12 | 2.4E-03 |
| TcCLB.510943.50 | delta-1-pyrroline-5-carboxylate dehydrogenase | 29.09 | 1.4E-04 |
| TcCLB.504105.140 | enolase | 19.27 | 2.5E-02 |
| TcCLB.509287.50 | glucose-6-phosphate 1-dehydrogenase | 8.08 | 1.0E-03 |
| TcCLB.509717.80 | glyceraldehyde 3-phosphate dehydrogenase, cytosolic | 20.63 | 8.2E-05 |
| TcCLB.506925.319 | isocitrate dehydrogenase | 11.92 | 3.1E-03 |
| TcCLB.511419.40 | phosphoglycerate kinase | 15.51 | 5.8E-05 |
| TcCLB.511281.60 | pyruvate kinase 2 | 8.50 | 1.7E-04 |
| TcCLB.507681.20 | succinyl-CoA ligase | 20.74 | 1.9E-02 |
| TcCLB.506355.50 | DnaJ chaperone protein | 30.43 | 4.5E-04 |
| TcCLB.511465.10 | proteasome activator protein pa26 | 20.92 | 3.2E-05 |
| TcCLB.506985.30 | proteasome alpha 3 subunit | 6.92 | 2.3E-03 |
| TcCLB.508043.30 | ubiquitin-activating enzyme E1 | 5.60 | 2.9E-03 |
| TcCLB.510661.19 | ubiquitin-activating enzyme E1 | 6.39 | 4.4E-03 |
| TcCLB.506583.30 | ubiquitin-like protein | 9.31 | 1.2E-03 |
| TcCLB.507029.30 | heat shock 70 kDa protein, mitochondrial precursor | 8.01 | 5.7E-04 |
| TcCLB.507831.60 | heat shock protein 110 | 22.22 | 3.6E-04 |
| TcCLB.508409.210 | Hsc70-interacting protein (Hip) | 9.16 | 9.1E-03 |

\* In orange is the TcZC3HTTP; in green, proteins related to RNA metabolism; in purple, metabolic proteins related to moonlighting functions; in yellow, proteins related to protein folding.

**Table 4.**
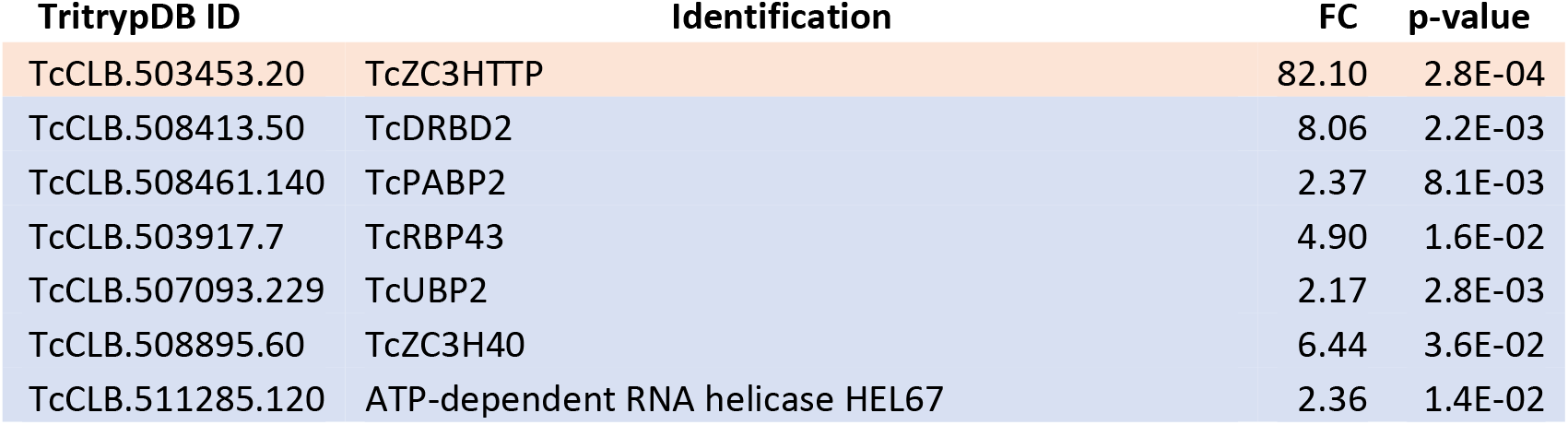
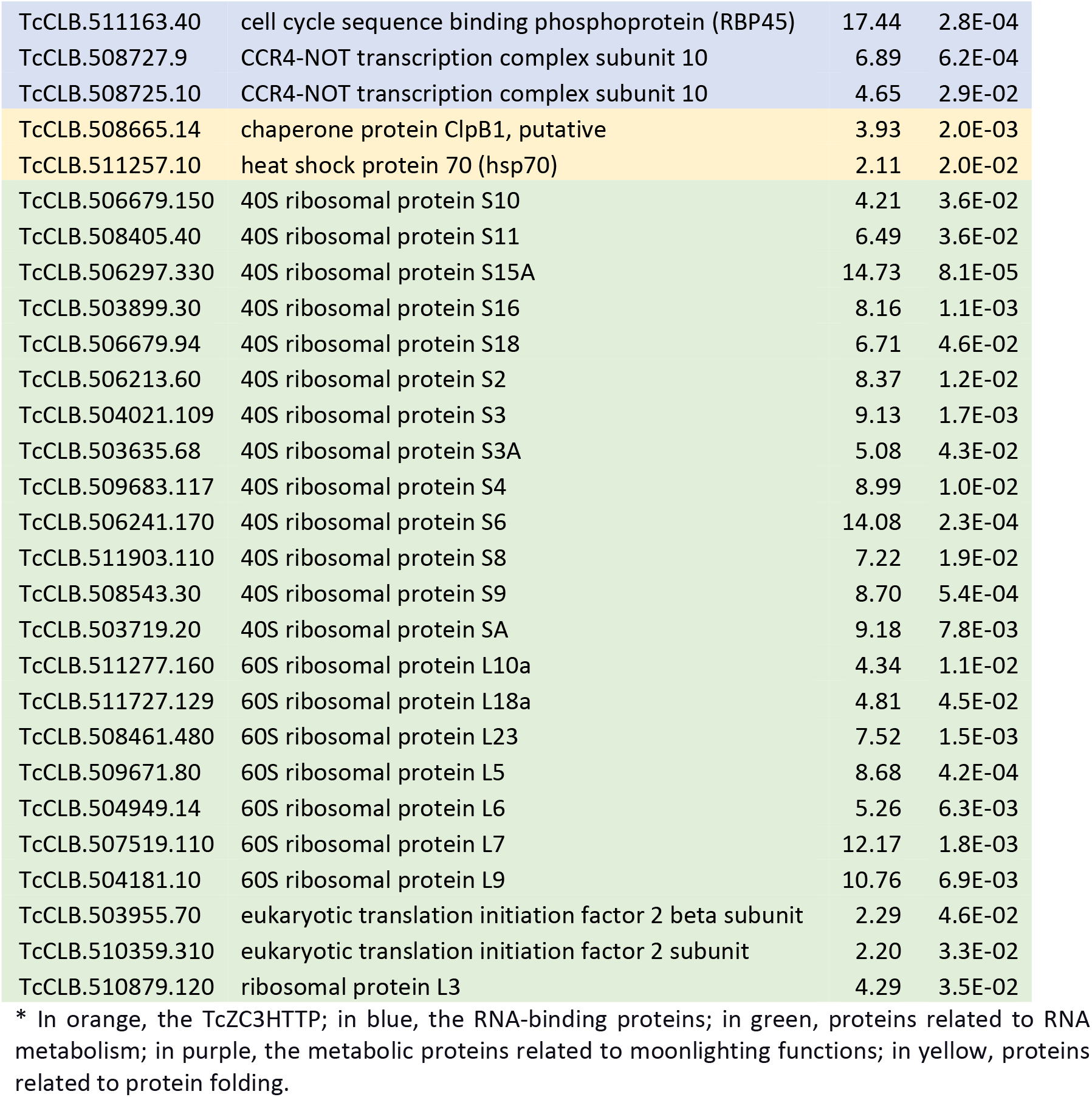
Proteins associated with TcZ3HTTP-3xFLAG in stressed epimastigotes by immunoprecipitation assay*.

| TritrypDB ID | Identification | FC | p-value |
| --- | --- | --- | --- |
| TcCLB.503453.20 | TcZC3HTTP | 82.10 | 2.8E-04 |
| TcCLB.508413.50 | TcDRBD2 | 8.06 | 2.2E-03 |
| TcCLB.508461.140 | TcPABP2 | 2.37 | 8.1E-03 |
| TcCLB.503917.7 | TcRBP43 | 4.90 | 1.6E-02 |
| TcCLB.507093.229 | TcUBP2 | 2.17 | 2.8E-03 |
| TcCLB.508895.60 | TcZC3H40 | 6.44 | 3.6E-02 |
| TcCLB.511285.120 | ATP-dependent RNA helicase HEL67 | 2.36 | 1.4E-02 |
| TcCLB.511163.40 | cell cycle sequence binding phosphoprotein (RBP45) | 17.44 | 2.8E-04 |
| TcCLB.508727.9 | CCR4-NOT transcription complex subunit 10 | 6.89 | 6.2E-04 |
| TcCLB.508725.10 | CCR4-NOT transcription complex subunit 10 | 4.65 | 2.9E-02 |
| TcCLB.508665.14 | chaperone protein ClpB1, putative | 3.93 | 2.0E-03 |
| TcCLB.511257.10 | heat shock protein 70 (hsp70) | 2.11 | 2.0E-02 |
| TcCLB.506679.150 | 40S ribosomal protein S10 | 4.21 | 3.6E-02 |
| TcCLB.508405.40 | 40S ribosomal protein S11 | 6.49 | 3.6E-02 |
| TcCLB.506297.330 | 40S ribosomal protein S15A | 14.73 | 8.1E-05 |
| TcCLB.503899.30 | 40S ribosomal protein S16 | 8.16 | 1.1E-03 |
| TcCLB.506679.94 | 40S ribosomal protein S18 | 6.71 | 4.6E-02 |
| TcCLB.506213.60 | 40S ribosomal protein S2 | 8.37 | 1.2E-02 |
| TcCLB.504021.109 | 40S ribosomal protein S3 | 9.13 | 1.7E-03 |
| TcCLB.503635.68 | 40S ribosomal protein S3A | 5.08 | 4.3E-02 |
| TcCLB.509683.117 | 40S ribosomal protein S4 | 8.99 | 1.0E-02 |
| TcCLB.506241.170 | 40S ribosomal protein S6 | 14.08 | 2.3E-04 |
| TcCLB.511903.110 | 40S ribosomal protein S8 | 7.22 | 1.9E-02 |
| TcCLB.508543.30 | 40S ribosomal protein S9 | 8.70 | 5.4E-04 |
| TcCLB.503719.20 | 40S ribosomal protein SA | 9.18 | 7.8E-03 |
| TcCLB.511277.160 | 60S ribosomal protein L10a | 4.34 | 1.1E-02 |
| TcCLB.511727.129 | 60S ribosomal protein L18a | 4.81 | 4.5E-02 |
| TcCLB.508461.480 | 60S ribosomal protein L23 | 7.52 | 1.5E-03 |
| TcCLB.509671.80 | 60S ribosomal protein L5 | 8.68 | 4.2E-04 |
| TcCLB.504949.14 | 60S ribosomal protein L6 | 5.26 | 6.3E-03 |
| TcCLB.507519.110 | 60S ribosomal protein L7 | 12.17 | 1.8E-03 |
| TcCLB.504181.10 | 60S ribosomal protein L9 | 10.76 | 6.9E-03 |
| TcCLB.503955.70 | eukaryotic translation initiation factor 2 beta subunit | 2.29 | 4.6E-02 |
| TcCLB.510359.310 | eukaryotic translation initiation factor 2 subunit | 2.20 | 3.3E-02 |
| TcCLB.510879.120 | ribosomal protein L3 | 4.29 | 3.5E-02 |

\* In orange, the TcZC3HTTP; in blue, the RNA-binding proteins; in green, proteins related to RNA metabolism; in purple, the metabolic proteins related to moonlighting functions; in yellow, proteins related to protein folding.

Ribosomal subunits 40S and 60S protein enrichment was observed in nutritionally stressed parasites. RBPs were also associated with TcZC3HTTP in this condition, including the zinc finger protein TcZC3H40, as well as the RRMs TcUPB2, TcDRBD2, TcRBP43, and TcPABP2 (Table 4). This result is in accordance with the RNAs associated with TcZC3HTTP in stress conditions. Since TcZC3HTTP was associated with clearly distinct protein groups under normal and nutritional stress, it is reasonable to assume that this RBP can perform different roles according to the cellular context. Furthermore, TcZC3HTTP association with other RBPs and ribosomal proteins during stress, instead of enzymes, implies its participation in the regulatory network rearrangement related to the stress response and is likely associated with a change in translation profiles.

### TcZC3HTTP disruption impact in *T. cruzi* gene expression (RNA)

One of the primary objectives of this study was to investigate the impact of TcZC3HTTP disruption on epimastigotes and nutritionally stressed cells by analyzing their transcriptome profiles. Our findings revealed differential expression of 126 messenger RNAs (mRNAs) in epimastigotes, with 96 mRNAs upregulated and 26 mRNAs downregulated in the absence of TcZC3HTTP (Figure 5 and Table S5).

**Figure 5.**
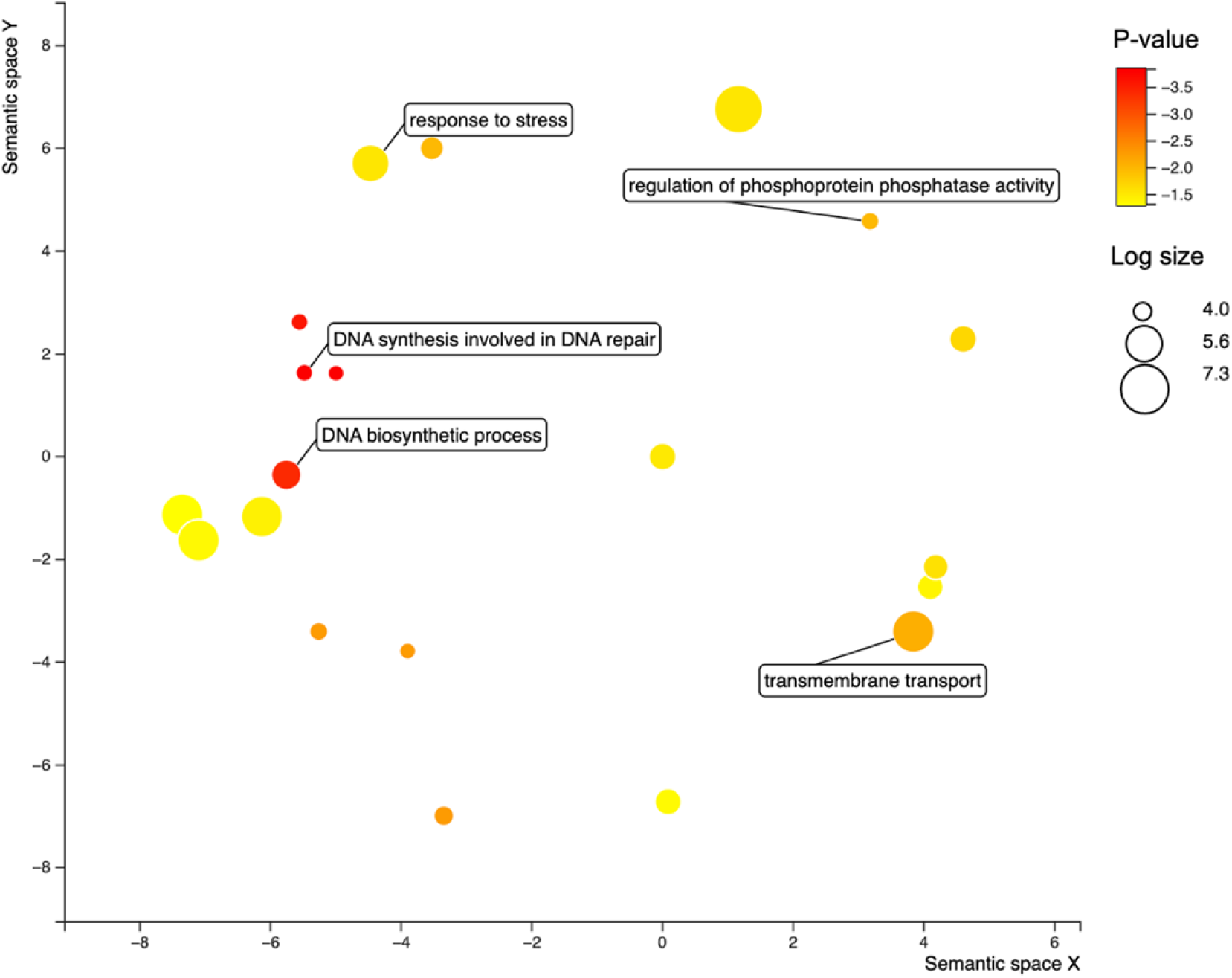
Bubble chart with the gene ontology terms enriched in the RNAs upregulated in the ΔTcZC3HTTP strain compared to the wild type. The colors refer to the p-value (log-10), and the bubble size the term counts (in log scale). The distance of the bubbles in the x- and y-axis refer to the proximity of the terms that belong to close-related biological processes.

In epimastigotes lacking TcZC3HTTP, an increase in the expression of mRNAs encoding proteins related to stress response, transmembrane transport, and DNA synthesis and repair was observed (Figure 5). The downregulated mRNAs were primarily related to surface proteins and proteins with predicted transmembrane domains. The results also indicated the downregulation of a chaperone DNAJ, several enzymes (including a kinase, alcohol dehydrogenase, metallopeptidase, exonuclease, and a dioxygenase), as well as the TcZC3HTTP transcript itself (as illustrated in Figure 5).

Interestingly, the TcZC3HTTP null mutation had minimal impact on the overall mRNA abundance of nutritionally stressed parasites. Only six transcripts were found to be differentially expressed, as shown in Table 3. The only upregulated transcript encodes a hypothetical protein with an unknown function domain (DUF229). The downregulated transcripts were an rRNA small subunit, an ubiquitin/ribosomal protein, and another hypothetical protein with no predicted domains. The other two transcripts that were less expressed were related to a protein known as PAD-8, which is associated with differentiation, and were also observed to be downregulated under both normal and stressed conditions (refer to Supplementary Table S5 and Table 3).

In our investigation, we sought to establish a correlation between the expression values of transcripts associated with TcZC3HTTP in the immunoprecipitation assay and the transcriptome of the knockout strain. Strikingly, the absence of TcZC3HTTP resulted in a noticeable downregulation or reduced expression levels of most of the associated transcripts, implying a potential role for this protein in stabilizing its target transcripts (Figure 6). This finding underscores the significance of TcZC3HTTP in post-transcriptional regulation and highlights its possible involvement in transcript stability.

**Figure 6.**
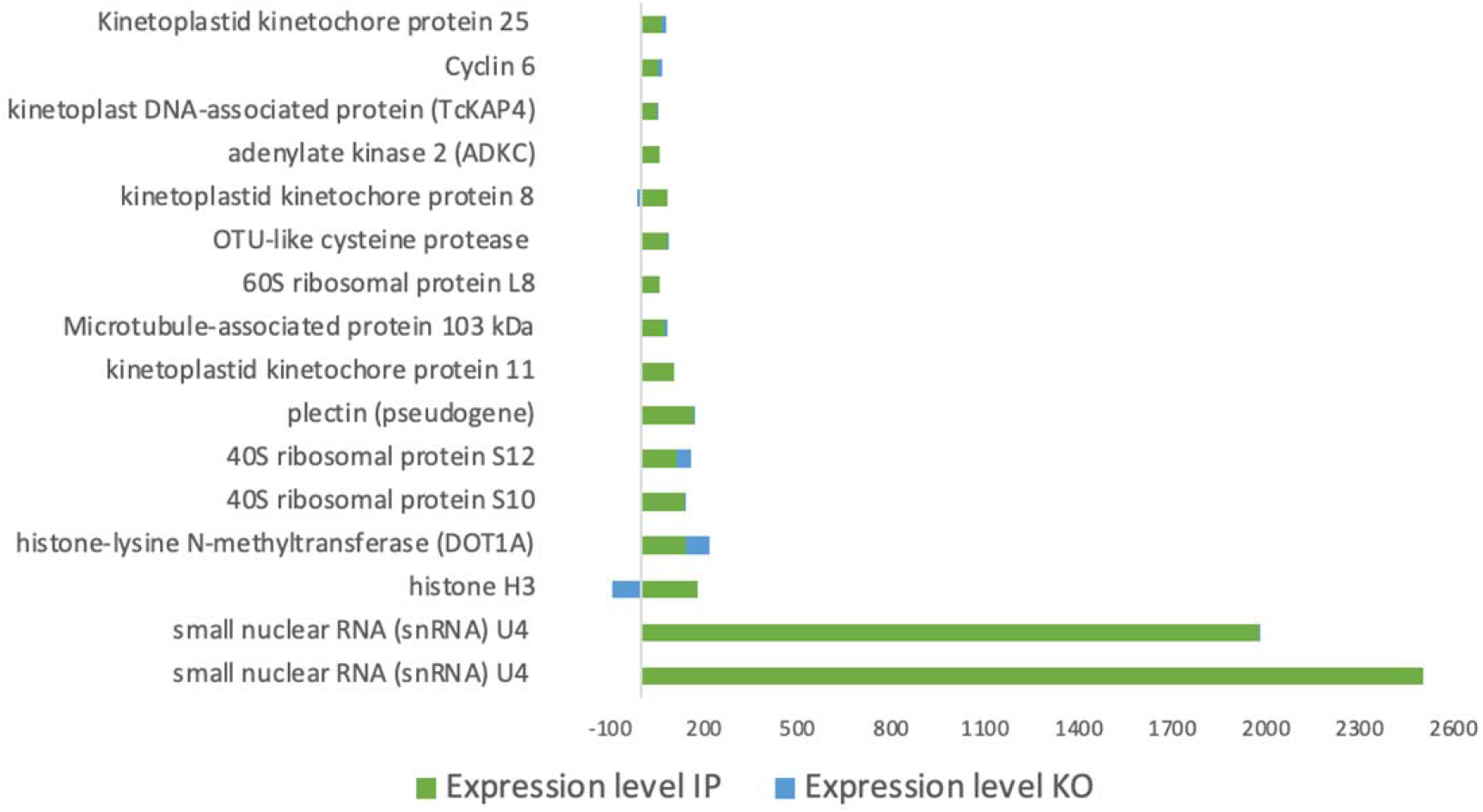
Bar chart plotted with the expression values of the transcripts enriched in the TcZC3HTTP immunoprecipitation (IP) assay (green) and the comparison with the expression values in the ΔTcZC3HTTP strain (Blue). The x-axis represents the expression values in TPM (transcripts per million).

### TcZC3HTTP disruption impact in *T. cruzi* gene expression regulation (Protein)

We conducted a proteomic analysis under normal and nutritional stress conditions to investigate the impact of TcZC3HTTP absence on *T. cruzi* gene expression. Our results revealed significant alterations in 65 proteins (32 upregulated and 33 downregulated) in epimastigotes, while 59 proteins (39 upregulated and 20 downregulated) exhibited differential expression in nutritionally stressed parasites (Figure 7 and Supplementary Tables S6 and S7).

**Figure 7.**
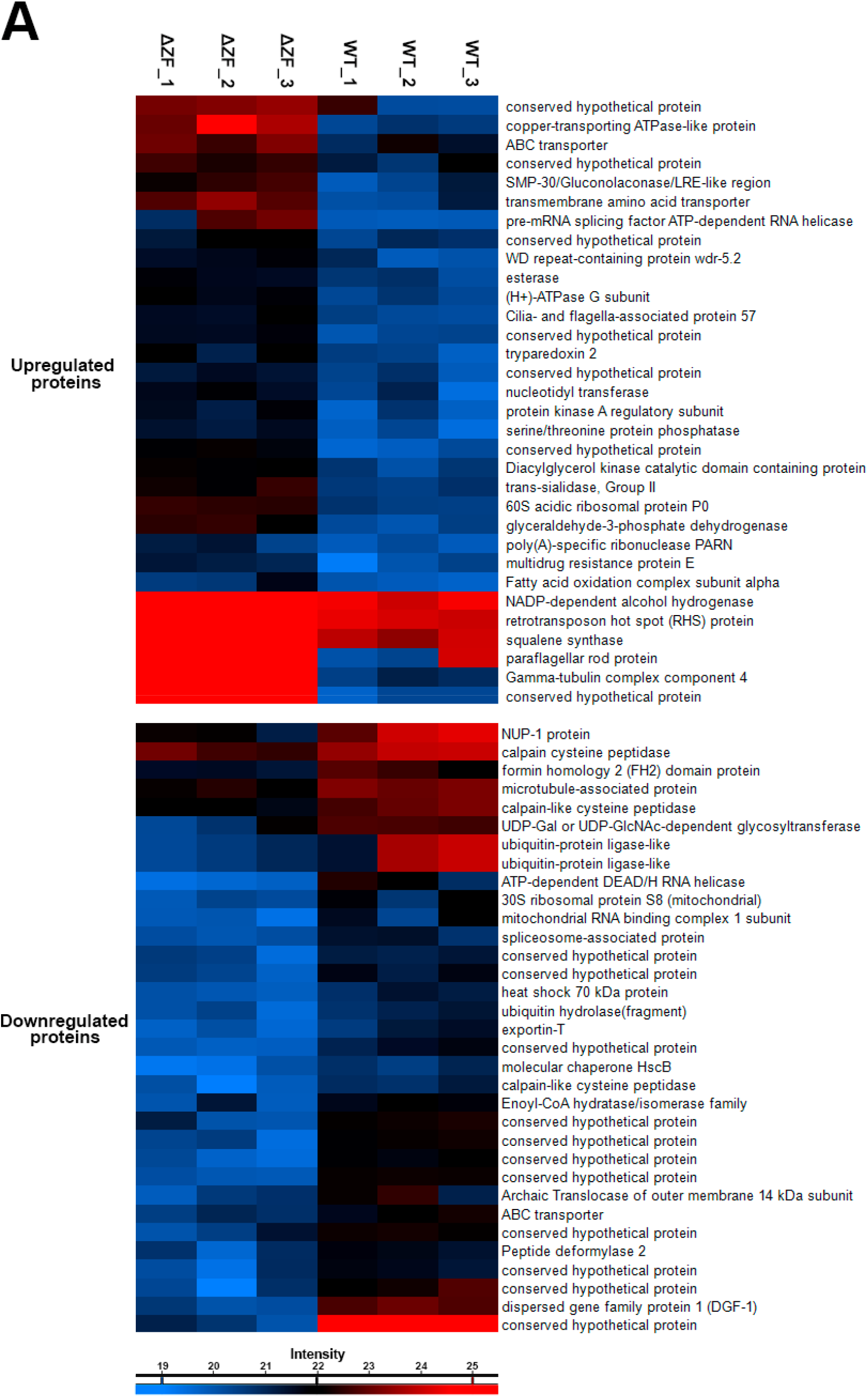
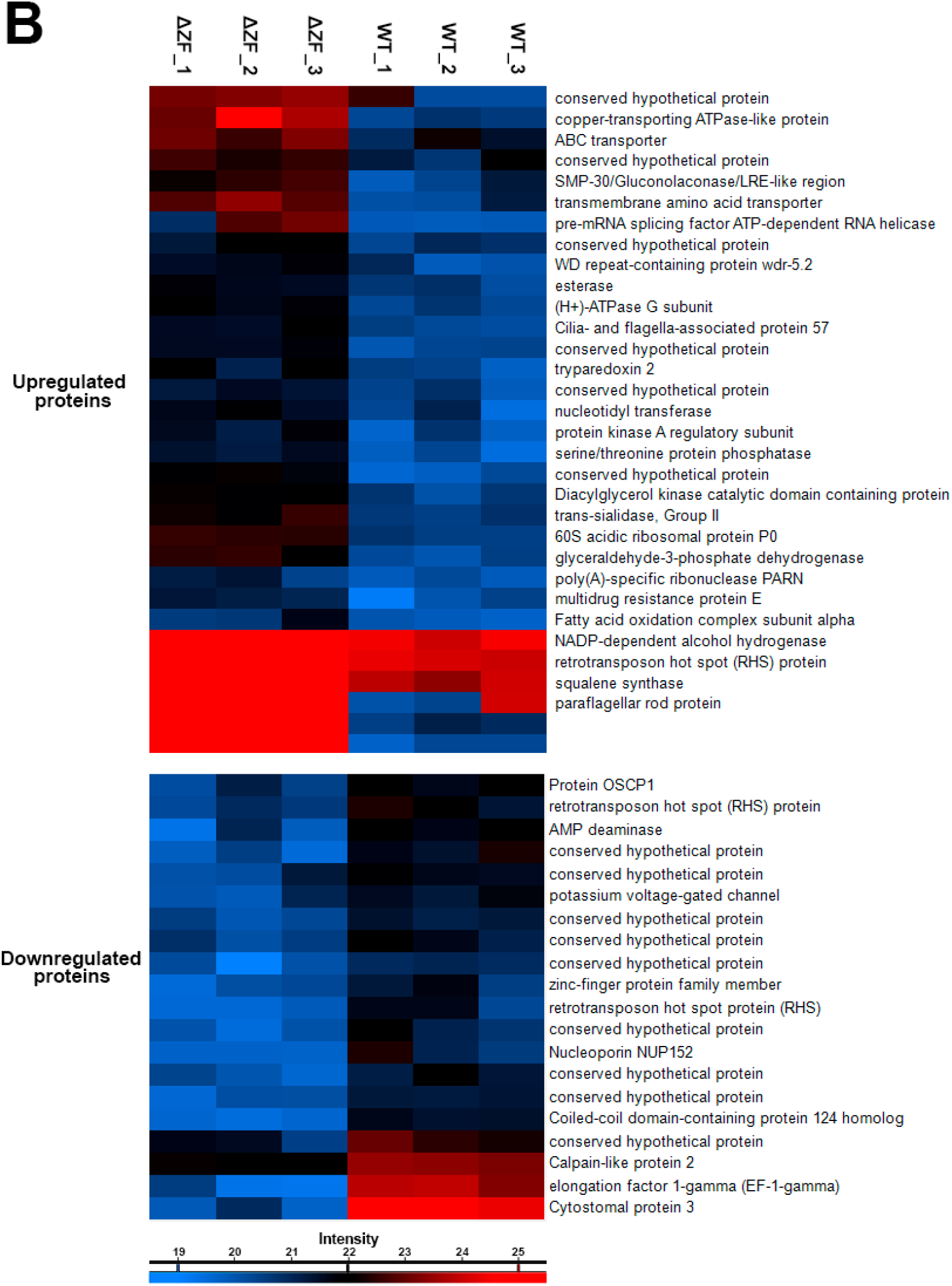
TcZC3HTTP disruption impact on epimastigotes and nutritionally stressed epimastigotes proteomic profile. Heat maps showing protein relative expression under normal growth (A) or during nutritional stress (B) epimastigotes lacking the TcZC3HTTP gene. In columns ΔZF_1, ΔZF_2, ΔZF_3, and WT_1, WT_2, and WT_3 are respective replicates for the TcZC3HTTP null mutant (ΔTcZC3HTTP) and wild-type populations. Identified protein expression levels are presented according to the colors from higher (red) to lower (blue) intensities, as indicated in the bars below each heat map set.

In epimastigotes lacking TcZC3HTTP, the upregulated proteins included a component of the gamma-tubulin complex 4, a paraflagellar rod protein, a copper-transporting ATPase-like protein, and a transmembrane amino acid transporter (Figure 7A). Among the downregulated proteins in normal conditions, we identified two ubiquitin-protein ligases, chaperones, an ATP-dependent DEAD/H RNA helicase, and three calpain-like cysteine peptidases. Furthermore, an exportin and a spliceosome-associated protein were detected (Figure 7A).

In nutritionally stressed parasites lacking TcZC3HTTP, we observed an upregulation of RBPs, ribosomal proteins, trans-sialidases (group II), and a rad21 double-strand-break repair protein homolog (Figure 7B). Notably, three proteins exhibited fold changes greater than 10, including the eukaryotic translation initiation factor 3, a conserved hypothetical protein with a predicted rhomboid-like domain, and a protein containing a tetratricopeptide-like helix and a SET domain. Among the 20 downregulated proteins in stressed parasites without TcZC3HTTP, we identified nucleoporin 149, a calpain-like protein, and an AMP deaminase (Figure 7B). The Cystostomal protein 3 and TcEIF1-γ were also downregulated, showing the most significant decreases in expression with fold changes of less than -20 and -15, respectively.

## DISCUSSION

Recently, we employed the CRISPR/Cas9 genome editing system to disrupt TcZC3HTTP and generated null mutants in *T. cruzi* (22). In addition to the TcZC3HTTP null mutant population, we also utilized populations expressing a tagged version of TcZC3HTTP to gain further insight into its role in *T. cruzi* gene expression regulation.

As previously described, numerous RBPs have been implicated in and shown to influence critical cellular processes, primarily through the modulation of their target transcripts (11–17). For instance, in *T. brucei*, disruption of the RBP TbRRM1 leads to cell cycle arrest, likely mediated by the modulation of its target TbNOP86, a nucleolar protein crucial for mitotic progression. Downregulation of TbNOP86 results in cell accumulation in the G2/M phase (15). Similarly, in *T. cruzi*, knockout of TcZC3H12 impairs cell proliferation, an effect linked to the targeting and stabilization of mRNAs encoding amino acid transporters (18). When examining the potential target mRNAs of TcZC3HTTP in epimastigotes, we observed an enrichment of transcripts encoding proteins involved in cell growth and division, such as cyclins, centrosomal proteins, and kinetochore proteins. Therefore, it is plausible that the contribution of TcZC3HTTP to cell proliferation is also mediated through its modulation of target transcripts.

Moonlighting proteins exhibit multiple functional roles, including enzymatic activity and RNA-binding capabilities, showcasing their versatility in cellular processes. Aconitase and glyceraldehyde-3-phosphate dehydrogenase (GAPDH) are prime examples of moonlighting proteins with RNA-binding properties (23–25). Aconitase acts as an enzyme in citrate interconversion and an iron regulatory protein (IRP1) by binding specific transcripts during iron deficiency (26). GAPDH, known for its glycolytic function, also binds RNA, though the underlying mechanism is not fully understood (27, 28). Other metabolic enzymes such as aldolase, lactate dehydrogenase, and glutamate dehydrogenase have been implicated in RNA interactions (25, 29). The expanding landscape of RNA-binding proteins encompasses structured RBPs, moonlighting proteins, and unstructured multitasking proteins. Notably, aldolase, trifunctional enzyme subunit b (HADHB), enolase 1 (ENO1), hydroxymethyltransferase (SHMT1), and pyruvate kinase M2 (PKM2) serve as further instances of RNA-binding metabolic enzymes, illuminating their moonlighting roles in RNA regulation (25, 30, 31). These findings provide valuable insights into the potential interplay between gene expression and intermediary metabolism orchestrated by these versatile RNA-binding metabolic enzymes.

We further identified the RBPs TcPUF9 and TcPUF5 transcripts immunoprecipitating with TcZC3HTTP in epimastigotes. TcPUF9 acts to stabilize its targets, and it seems that this occurs specifically in the cell cycle S-phase (11, 32). RBP45 is a cell cycle sequence binding phosphoprotein involved in cell cycle regulation. In *T. brucei*, it is associated with the regulation of transcripts during the S phase. RBP45, TcZC3H39, and TcZC3H40, and their respective orthologs CSBPA and CSBPB, interact and form a complex with RBP33. Although RBP33 was not identified in our initial analyses, the mRNA of RBP45 and the protein co-immunoprecipitated with TcZC3HTTP. This is an important observation, especially considering that TcZC3H40 was detected in the immunoprecipitation of TcZC3HTTP under stress conditions.

During stress conditions, TcZC3HTTP was found to associate with ribosomal RNAs, mRNAs encoding tubulins, kinases, and several other RBPs, including TcZC3H11, TcZC3H35, TcZC3H28, and TcPUF6. It is worth noting that TcPABP2 mRNA exhibited a high enrichment under stress conditions (fold change > 15), indicating its potential involvement in stress response, translation profile changes, and rearrangements of RNA granules, where RBPs play a crucial role and coordinate their actions. These findings further support the concept of the cytoskeleton as a regulon, which is consistent with the similarities observed with TcZC3H39 (33).

TcZC3HTTP binds to its mRNA in both studied conditions. This observation has been reported in *T. cruzi* for the zinc finger protein 2 (TcZFP2), which interacts with its mRNA through an A-rich sequence in the 3’ UTR (34). Similarly, tcrbp19 mRNA was identified as a target of its protein, TcRBP19, through its 3’ UTR. However, in this case, TcRBP19 is responsible for negatively regulating its own mRNA rather than stabilizing it (35).

In conclusion, our study generated TcZC3HTTP null mutants in *T. cruzi* and investigated its role in gene expression regulation. We found that TcZC3HTTP potentially modulates target transcripts associated with cell growth and division. Immunoprecipitation assays demonstrated the interaction of TcZC3HTTP with other RBPs, such as TcPUF9, further emphasizing its involvement in RNA regulation. Interestingly, TcZC3HTTP exhibited enrichment with specific mRNAs during stress conditions, suggesting its role in stress response and mRNP rearrangements. Overall, our study provides valuable insights into the multifaceted roles of TcZC3HTTP in *T. cruzi* and its contribution to cellular processes and stress responses.

## METHODS

### Phylogenetic Analysis

The ZC3HTTP sequences from Trypanosomatids were obtained by using *T. cruzi* protein (TCDM_03704, ESS67632.1) sequence as a query on BLASTp (36, 37) searches on TritrypDB (38) and GenBank (39) databases. The MAFFT online tool aligned the sequences (40), and poorly aligned positions were discarded by TrimAl (41). The best model with computation of rates and support was determined by PhyML analysis using Phylemon 2.0 web suite (42). The consensus phylogenetic tree was visualized and edited using FigTree v1.4.4 program (available at http://tree.bio.ed.ac.uk/software/figtree).

### *Trypanosoma cruzi* culture and TcZC3HTTP transfected and null mutant parasites

*T. cruzi* Dm28c epimastigotes were cultured at 28 °C in liver infusion tryptose (LIT) medium supplemented with 10% heat-inactivated fetal bovine serum (FBS). Cultures overexpressing or null mutants for TcZC3HTTP (TCDM_03704) were obtained as previously described (22). Briefly, 5.10^6^ early-log phase wild-type epimastigotes were transfected with 50 µg of pTcGW plasmid (43) containing the gene *tczc3http* (pTcGW-TcZC3HTTP) to generate a TcZC3HTTP-3xFLAG population, selected with G418 (500 µg.ml-1) treatment. TcZC3HTTP null mutant parasites were generated from *T. cruzi* Cas9-expressing lineages (for more details, see (21)) transfected with gRNA targeting the gene tczc3http along with a respective DNA donor to insert a sequence encoding stop codons in three different frames into the target gene. Transfectant and null mutant parasite confirmation are shown elsewhere (22).

### Nutritional stress, cell proliferation, and cell cycle analysis

Nutritional stress was performed as previously described (33). Briefly, epimastigotes at the end of the exponential growth phase (five-day cultures at a density ≥ 5.10^7^ parasites.ml-1) were harvested by centrifugation at 5,000 x g for 5 min at room temperature, washed twice with PBS (pH 7.4), and incubated for 2 h at 28 °C in TAU medium (190 mM NaCl, 17 mM KCl, 2 mM MgCl2, 2 mM CaCl2, 8 mM phosphate buffer pH 6.0) at a density of 5.10^8^ parasites.ml-1. For cell proliferation assay, cultures starting with 1.10^6^ epimastigotes.ml-1 were monitored throughout 7 days, and parasite density was determined every 24 – 48 h using a hemocytometer. The cell cycle was assessed by flow cytometry as previously described (21). After DNA content determination, cell cycle phases were defined using the model Watson Pragmatic algorithm (44) in the FlowJoTM v10.8.1 software. All experiments were performed at least in technical and biological triplicates, and One-way ANOVA statically analyzed the data. Graphs and statistics were obtained using the GraphPad Prism software version 8.4.2 (La Jolla, CA, USA). The immunofluorescence assay was performed as previously described (22).

### RNA and protein enrichment by immunoprecipitation

To identify potential TcZC3HTTP RNAs targets and protein partners, proteins and RNAs were enriched by immunoprecipitation using magnetic beads coupled with anti-FLAG antibodies. 10^9^ parasites expressing TcZC3HTTP-FLAG (in log phase or nutritionally stressed in TAU pH 6.0 for 2 h) were harvested by centrifugation (3000 x g for 5 min), washed with PBS (TAU for stress condition), and lysed accordingly to extract and capture RNAs or proteins. For RNAs, parasites were incubated in lysis buffer (NaCl 150mM, 20mM Tris-HCl pH 7.4, and Nonidet® P40 0.5 %) at 4 °C for 10 min (lysis was confirmed by optical microscopy). The cellular extract was centrifuged (10000 x g for 20 min at 4 °C) and incubated with anti-FLAG M2 magnetic beads (Sigma) for 2h at 4 °C following washing and elution steps using a magnetic separation stand according to the manufacturer instructions. RNAs from the eluted fraction were extracted using de miRCURY RNA Isolation Kit Cell & Plant (QIAGEN) and stored at -80 °C until cDNA library preparation. For protein isolation and enrichment, cells were incubated with a cold cavitation buffer (20 mM HEPES KOH pH 7.4, 75 mM potassium acetate, 4 mM magnesium acetate, and 2 mM DTT) supplemented with 1 x cOmplete ™ Protease Inhibitor Cocktail (Sigma) and 40 µg.ml-1 RNase A (Sigma) and disrupted using 1000psi (70 bar) for 40 min at 4 °C. Next, the lysate was centrifuged (10000 x g for 10 min at 4 °C). The supernatant was incubated with magnetic beads Dynabeads ™ M-280 Sheep anti-mouse IgG (Sigma) conjugated with monoclonal ANTI-FLAG ® M2 antibody (Sigma) for 2h at 4 °C. Later, the magnetic beads were washed, and proteins were eluted in Laemmli sample buffer and stored at -80 °C.

### TcZC3HTTP null mutant RNAs and protein extraction

To investigate the impact of TcZC3HTTP absence in *T. cruzi* RNAs and proteins, total RNA and protein were extracted from TcZC3HTTP null mutant epimastigotes in both log phase and nutritionally stressed conditions. Total RNA was obtained from 10^8^ parasites that were harvested (3000 x g for 5 min), washed with PBS (TAU for stress conditions), and lysed using the RNeasy Plus micro kit (QIAGEN) following fabricant instructions. Then, the extracted RNAs were stored at -80 °C. For protein extraction, 3.10^7^ parasites were used, and after harvesting and washing steps, the cells were lysed in a Laemmli sample buffer and stored at -80 °C.

### Transcriptomic analysis

Before proceeding to cDNA library preparation, all RNA samples were quantified, and their integrity was checked. For RNA quantification, Qubit Fluorometric Quantitation (Thermo Fisher Scientific) RNA BR kit was used. The fragment distribution, integrity, and quality analysis were performed on a 2100 Bioanalyzer (Agilent Technologies, Agilent RNA 6000 Pico Kit). According to the manufacturer’s instructions, the Illumina TruSeq Stranded mRNA kit (Illumina) was used to build the null mutant and IP libraries. The library concentration was measured with Qubit 4 (Thermo Fisher Scientific) DNA HS assay (Invitrogen), and 2100 Bioanalyzer DNA 1000 kit (Agilent) was used to determine the medium peak size. Sequencing was performed on a MiSeq (Illumina, v2 pair-end 75-bp run) sequencer at Carlos Chagas Institute (Fiocruz Paraná). Each sample was individually added to the two lanes as a technical replica to avoid lane-specific errors and improve coverage. Data analysis was performed with CLC Genomics Workbench v 20.0.03 software (Qiagen). First, each sample was checked for quality and filtered above PHRED Q30. Reads were mapped against the reference *T. cruzi* genome (CL Brener strain GCA_000209065.1 ASM20906v1) following the parameters: match score (1), mismatch cost (2), linear insertion cost (3), deletion cost (3), length fraction (0.6), similarity fraction (0.8), and global alignment. Differential expression analysis was applied using the mapped read counts. Transcripts were considered differentially expressed if presenting fold change equal or superior to 3 in the immunoprecipitation assays and 1.5 in total RNA identification. A false discovery rate (FDR) ≤ 0.05 was also used. For gene ontology analysis, we used the DAVID functional annotation tool (45).

### Proteomic analysis

Proteins extracted by immunoprecipitation were analyzed by electrophoresis followed by silver staining and western blot to confirm the presence and enrichment of TcZC3HTTP-3xFLAG. Then, the total protein extracts (approximately 100ug) and the proteins extracted by immunoprecipitation were separated in 13% polyacrylamide gels. The lanes were reduced with DTT, alkylated, and digested with trypsin. The peptides were extracted with acetonitrile and trifluoroacetic acid, dried, and desalted. Peptides of each sample were separated by online reversed-phase nanoscale capillary liquid chromatography (Ultimate 3000 RSLCnano chromatograph) and analyzed by nano-electrospray mass spectrometry. The mass spectrometry assay was performed at the facility RPT02H of Carlos Chagas Institute (Fiocruz, Parana) Orbitrap Fusion Lumos mass spectrometer. The mass spectrometry data was matched against the *T. cruzi* Dm28c GenBank database (GCA_003177105.1) with MaxQuant software v2.0.3.0 with quantification set to the LFQ method and the following parameters: cysteine carbamidomethylation set to fixed modification, methionine oxidation and N-terminal acetylation set to variable modification, an FDR of 1% for peptide and protein identification was independently applied. The resulting peptides were analyzed with Perseus v.1.6.14.0. Contaminants, reverse sequences, and peptides only identified by site were removed, the LFQ intensities were transformed to log2(x) scale, and a normal distribution imputed missing value. T-tests were performed to determine differentially expressed proteins (p-value ≤ 0.05) and fold change ≥ 2.

## Data analysis

The RNA-seq data have been deposited at the Sequence Read Archive (SRA) database under the accession number (SRA: PRJNA986951).

## ACKNOWLEDGMENTS

Fiocruz, grant number ICC-008-FIO-21-2-14, funded LRA and SG. SG is a research fellow awardee from CNPq.

## Author Contributions

BAAR performed most of the experiment and wrote the manuscript. LRA performed the transcriptomic analysis and wrote the manuscript. ACL and IFMR performed the proteomic analysis. ERF performed the phylogenetic analysis. MLR and SG discussed all the results and contributed to writing the manuscript. All the authors approved the final version of the manuscript.

## Supplementary Tables

Supplemental Table 1 – RNA-seq of the immunoprecipitated transcripts associated with TcZC3HTTP in epimastigote.

Supplemental Table 2 - RNA-seq of the immunoprecipitated transcripts associated with TcZC3HTTP in stressed epimastigote.

Supplemental Table 3 – Proteins associated with TcTZC3HTTP in epimastigote.

Supplemental Table 4 - Proteins associated with TcTZC3HTTP under stress conditions.

Supplemental Table 5 – Transcriptome of ΔTcTZC3HTTP compared with the wild-type strain in epimastigote.

Supplemental Table 6 - Proteome of ΔTcTZC3HTTP compared with the wild-type strain in epimastigote.

Supplemental Table 7- Proteome of ΔTcTZC3HTTP compared with the wild-type strain in stressed epimastigote.

---

**Supplementary Figure 1.**
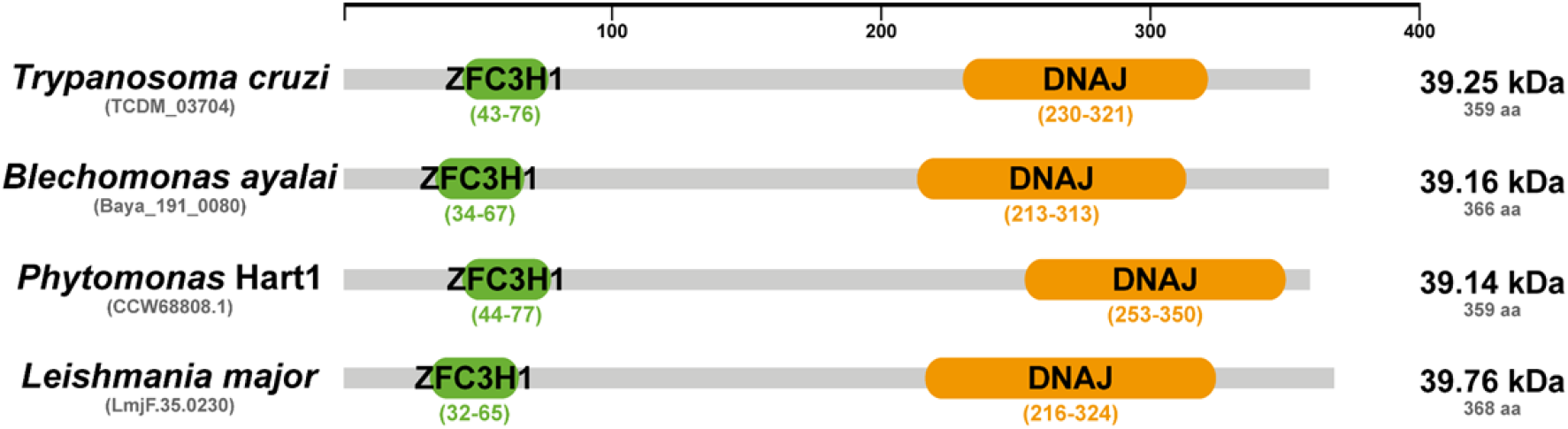
Schematic representation of ZC3HTTP homologs from four Trypanosomatids species. The proteins ZC3HTTP from *T. cruzi, B. ayalai*, Phytomonas Hart1, and *L. major* show similar size and relative positioning for their functional domains, revealing high conservation between groups within trypanosomatids. The full-length proteins are represented in gray, while Zinc finger ZFC3H1 domains are represented in green, and Chaperone DNAJ domains are represented in orange. The domain boundaries were generated through sequence searches using the Interpro online tool (https://www.ebi.ac.uk/interpro/search/sequence/)

## References

1. Clayton CE. 2002. Life without transcriptional control? From fly to man and back again. EMBO J.

2. Martínez-Calvillo S, Vizuet-de-Rueda JC, Florencio-Martínez LE, Manning-Cela RG, Figueroa-Angulo EE. 2010. Gene expression in trypanosomatid parasites. J Biomed Biotechnol.

3. El-Sayed NM, Myler PJ, Bartholomeu DC, Nilsson D, Aggarwal G, Tran A-N, Ghedin E, Worthey EA, Delcher AL, Blandin G, Westenberger SJ, Caler E, Cerqueira GC, Branche C, Haas B, Anupama A, Arner E, Aslund L, Attipoe P, Bontempi E, Bringaud F, Burton P, Cadag E, Campbell DA, Carrington M, Crabtree J, Darban H, da Silveira JF, de Jong P, Edwards K, Englund PT, Fazelina G, Feldblyum T, Ferella M, Frasch AC, Gull K, Horn D, Hou L, Huang Y, Kindlund E, Klingbeil M, Kluge S, Koo H, Lacerda D, Levin MJ, Lorenzi H, Louie T, Machado CR, McCulloch R, McKenna A, Mizuno Y, Mottram JC, Nelson S, Ochaya S, Osoegawa K, Pai G, Parsons M, Pentony M, Pettersson U, Pop M, Ramirez JL, Rinta J, Robertson L, Salzberg SL, Sanchez DO, Seyler A, Sharma R, Shetty J, Simpson AJ, Sisk E, Tammi MT, Tarleton R, Teixeira S, Van Aken S, Vogt C, Ward PN, Wickstead B, Wortman J, White O, Fraser CM, Stuart KD, Andersson B. 2005. The genome sequence of Trypanosoma cruzi, etiologic agent of Chagas disease. Science 309:409–415.

4. Herreros-Cabello A, Callejas-Hernández F, Gironès N, Fresno M. 2020. Trypanosoma Cruzi Genome: Organization, Multi-Gene Families, Transcription, and Biological Implications. Genes 11:1196.

5. Keene JD. 2007. RNA regulons: coordination of post-transcriptional events. Nat Rev Genet.

6. Glisovic T, Bachorik JL, Yong J, Dreyfuss G. 2008. RNA-binding proteins and post-transcriptional gene regulation. FEBS Lett.

7. Decker CJ, Parker R. 2012. P-bodies and stress granules: possible roles in the control of translation and mRNA degradation. Cold Spring Harb Perspect Biol.

8. Alves LR. 2016. RNA-binding proteins related to stress response and differentiation in protozoa. World Journal of Biological Chemistry 7:78.

9. Kolev NG, Ullu E, Tschudi C. 2014. The emerging role of RNA-binding proteins in the life cycle of Trypanosoma brucei. Cell Microbiol.

10. Lueong S, Merce C, Fischer B, Hoheisel JD, Erben ED. 2016. Gene expression regulatory networks in Trypanosoma brucei: insights into the role of the mRNA-binding proteome. Molecular Microbiology 100:457–471.

11. Archer SK, Luu VD, de Queiroz RA, Brems S, Clayton C. 2009. Trypanosoma brucei PUF9 regulates mRNAs for proteins involved in replicative processes over the cell cycle. PLoS Pathog.

12. Das A, Morales R, Banday M, Garcia S, Hao L, Cross GA, Estevez AM, Bellofatto V. 2012. The essential polysome-associated RNA-binding protein RBP42 targets mRNAs involved in Trypanosoma brucei energy metabolism. RNA.

13. Wurst M, Seliger B, Jha BA, Klein C, Queiroz R, Clayton C. 2012. Expression of the RNA recognition motif protein RBP10 promotes a bloodstream-form transcript pattern in Trypanosoma brucei: Trypanosome RBP10 function. Molecular Microbiology 83:1048–1063.

14. Droll D, Minia I, Fadda A, Singh A, Stewart M, Queiroz R, Clayton C. 2013. Post-transcriptional regulation of the trypanosome heat shock response by a zinc finger protein. PLoS Pathog.

15. Levy GV, Bañuelos CP, Níttolo AG, Ortiz GE, Mendiondo N, Moretti G, Tekiel VS, Sánchez DO. 2015. Depletion of the SR-related protein TbRRM1 leads to cell cycle arrest and apoptosis-like death in Trypanosoma brucei. PLoS ONE 10:1–20.

16. Alcantara MV, Kessler RL, Gonçalves REG, Marliére NP, Guarneri AA, Picchi GFA, Fragoso SP. 2018. Knockout of the CCCH zinc finger protein TcZC3H31 blocks Trypanosoma cruzi differentiation into the infective metacyclic form. Molecular and Biochemical Parasitology https://doi.org/10.1016/j.molbiopara.2018.01.006.

17. Bishola Tshitenge T, Clayton C. 2022. The *Trypanosoma brucei* RNA-binding protein DRBD18 ensures correct mRNA *trans* splicing and polyadenylation patterns. RNA 28:1239–1262.

18. Tavares TS, Mügge FLB, Grazielle-Silva V, Valente BM, Goes WM, Oliveira AER, Belew AT, Guarneri AA, Pais FS, El-Sayed NM, Teixeira SMR. 2021. A *Trypanosoma cruzi* zinc finger protein that is implicated in the control of epimastigote-specific gene expression and metacyclogenesis. Parasitology 148:1171–1185.

19. Lander N, Li ZH, Niyogi S, Docampo R. 2015. CRISPR/Cas9-induced disruption of paraflagellar rod protein 1 and 2 genes in Trypanosoma cruzi reveals their role in flagellar attachment. mBio 6.

20. Burle-Caldas GA, Soares-Simões M, Lemos-Pechnicki L, DaRocha WD, Teixeira SMR. 2018. Assessment of two CRISPR-Cas9 genome editing protocols for rapid generation of Trypanosoma cruzi gene knockout mutants. International Journal for Parasitology 48:591–596.

21. Romagnoli BAA, Picchi GFA, Hiraiwa PM, Borges BS, Alves LR, Goldenberg S. 2018. Improvements in the CRISPR/Cas9 system for high efficiency gene disruption in Trypanosoma cruzi. Acta Tropica 178:190–195.

22. Romagnoli BAA, Holetz FB, Alves LR, Goldenberg S. 2020. RNA Binding Proteins and Gene Expression Regulation in Trypanosoma cruzi. Frontiers in Cellular and Infection Microbiology https://doi.org/10.3389/fcimb.2020.00056.

23. Jeffery CJ. 1999. Moonlighting proteins. Trends Biochem Sci 24:8–11.

24. Collingridge PW, Brown RW, Ginger ML. 2010. Moonlighting enzymes in parasitic protozoa. Parasitology.

25. Balcerak A, Trebinska-Stryjewska A, Konopinski R, Wakula M, Grzybowska EA. 2019. RNA– protein interactions: disorder, moonlighting and junk contribute to eukaryotic complexity. Open Biol 9:190096.

26. Leipuviene R, Theil EC. 2007. The family of iron responsive RNA structures regulated by changes in cellular iron and oxygen. Cell Mol Life Sci 64:2945–2955.

27. Tristan C, Shahani N, Sedlak TW, Sawa A. 2011. The diverse functions of GAPDH: Views from different subcellular compartments. Cellular Signalling 23:317–323.

28. White MR, Garcin ED. 2016. The sweet side of RNA regulation: glyceraldehyde-3-phosphate dehydrogenase as a noncanonical RNA -binding protein. WIREs RNA 7:53–70.

29. Castello A, Hentze MW, Preiss T. 2015. Metabolic Enzymes Enjoying New Partnerships as RNA-Binding Proteins. Trends in Endocrinology & Metabolism 26:746–757.

30. Castello A, Horos R, Strein C, Fischer B, Eichelbaum K, Steinmetz LM, Krijgsveld J, Hentze MW. 2013. System-wide identification of RNA-binding proteins by interactome capture. Nat Protoc 8:491–500.

31. Castello A, Hentze MW, Preiss T. 2015. Metabolic Enzymes Enjoying New Partnerships as RNA-Binding Proteins. Trends in Endocrinology & Metabolism 26:746–757.

32. Kramer S. 2012. Developmental regulation of gene expression in the absence of transcriptional control: The case of kinetoplastids Molecular and Biochemical Parasitology.

33. Alves LR, Oliveira C, Mörking PA, Kessler RL, Martins ST, Romagnoli BA, Marchini FK, Goldenberg S. 2014. The mRNAs associated to a zinc finger protein from Trypanosoma cruzi shift during stress conditions. RNA Biol.

34. Mörking PA, Rampazzo R e C, Walrad P, Probst CM, Soares MJ, Gradia DF, Pavoni DP, Krieger MA, Matthews K, Goldenberg S, Fragoso SP, Dallagiovanna B. 2012. The zinc finger protein TcZFP2 binds target mRNAs enriched during Trypanosoma cruzi metacyclogenesis. Mem Inst Oswaldo Cruz.

35. Pérez-Díaz L, Correa A, Moretão MP, Goldenberg S, Dallagiovanna B, Garat B. 2012. The overexpression of the trypanosomatid-exclusive TcRBP19 RNA-binding protein affects cellular infection by Trypanosoma cruzi. Mem Inst Oswaldo Cruz.

36. Altschul SF, Madden TL, Schäffer AA, Zhang J, Zhang Z, Miller W, Lipman DJ. 1997. Gapped BLAST and PSI-BLAST: a new generation of protein database search programs. Nucleic Acids Research 25:3389–3402.

37. Camacho C, Coulouris G, Avagyan V, Ma N, Papadopoulos J, Bealer K, Madden TL. 2009. BLAST+: architecture and applications. BMC Bioinformatics 10:421.

38. Aslett M, Aurrecoechea C, Berriman M, Brestelli J, Brunk BP, Carrington M, Depledge DP, Fischer S, Gajria B, Gao X, Gardner MJ, Gingle A, Grant G, Harb OS, Heiges M, Hertz-Fowler C, Houston R, Innamorato F, Iodice J, Kissinger JC, Kraemer E, Li W, Logan FJ, Miller JA, Mitra S, Myler PJ, Nayak V, Pennington C, Phan I, Pinney DF, Ramasamy G, Rogers MB, Roos DS, Ross C, Sivam D, Smith DF, Srinivasamoorthy G, Stoeckert CJ, Subramanian S, Thibodeau R, Tivey A, Treatman C, Velarde G, Wang H. 2009. TriTrypDB: A functional genomic resource for the Trypanosomatidae. Nucleic Acids Research 38.

39. Benson DA, Karsch-Mizrachi I, Lipman DJ, Ostell J, Wheeler DL. 2007. GenBank. Nucleic Acids Research 36:D25–D30.

40. Katoh K. 2002. MAFFT: a novel method for rapid multiple sequence alignment based on fast Fourier transform. Nucleic Acids Research 30:3059–3066.

41. Capella-Gutierrez S, Silla-Martinez JM, Gabaldon T. 2009. trimAl: a tool for automated alignment trimming in large-scale phylogenetic analyses. Bioinformatics 25:1972–1973.

42. Sanchez R, Serra F, Tarraga J, Medina I, Carbonell J, Pulido L, De Maria A, Capella-Gutierrez S, Huerta-Cepas J, Gabaldon T, Dopazo J, Dopazo H. 2011. Phylemon 2.0: a suite of web-tools for molecular evolution, phylogenetics, phylogenomics and hypotheses testing. Nucleic Acids Research 39:W470–W474.

43. Batista M, Marchini FK, Celedon PA, Fragoso SP, Probst CM, Preti H, Ozaki LS, Buck GA, Goldenberg S, Krieger MA. 2010. A high-throughput cloning system for reverse genetics in Trypanosoma cruzi. BMC Microbiology 10:259.

44. Watson JV, Chambers SH, Smith PJ. 1987. A pragmatic approach to the analysis of DNA histograms with a definable G1 peak. Cytometry 8:1–8.

45. Huang da W, Sherman BT, Lempicki RA. 2009. Systematic and integrative analysis of large gene lists using DAVID bioinformatics resources. Nat Protoc.

